# Mosaic heterochrony in neural progenitors sustains accelerated brain growth and neurogenesis in the juvenile killifish *N. furzeri*

**DOI:** 10.1101/747477

**Authors:** Marion Coolen, Miriam Labusch, Abdelkrim Mannioui, Beate Hoppe, Mario Baumgart, Laure Bally-Cuif

## Abstract

While developmental mechanisms driving increase in brain size during vertebrate evolution are actively studied, we know less of evolutionary strategies allowing to boost brain growth speed. In zebrafish and other vertebrates studied to date, radial glia (RG) constitute the primary neurogenic progenitor population throughout life (Kriegstein and Alvarez-Buylla, 2009); thus, RG activity is a determining factor of growth speed. Here, we ask whether enhanced RG activity is the mechanism selected to drive explosive growth, in adaptation to an ephemeral habitat. In post-hatching larvae of the turquoise killifish, which display drastic developmental acceleration, we show that the dorsal telencephalon (pallium) grows three times faster than in zebrafish. Rather than resulting from enhanced RG activity, we demonstrate that pallial growth is the product of a second type of progenitors (that we term AP for apical progenitors) that actively sustains neurogenesis and germinal zone self-renewal. Intriguingly, AP appear to retain, at larval stages, features of early embryonic progenitors. In parallel, RG enter premature quiescence and express markers of astroglial function. Together, we propose that mosaic heterochrony within the neural progenitor population may permit rapid pallial growth by safeguarding both continued neurogenesis and astroglial function.

## Results

### The killifish as a novel model to study progenitors dynamics driving accelerated brain growth

The dorsal part of the telencephalon, or pallium, varies substantially in size and organization between Vertebrate species, an evolutionary plasticity most likely linked to the crucial adaptive functions of high integrative centers hosted by this brain region. Many studies have focussed on developmental mechanisms contributing to the expansion of the pallium, at the origin of the mammalian cortex diversification (Lui et al., 2011; Romero and Borrell, 2015). In contrast, we know little about developmental mechanisms that would allow speeding up pallial development. Teleost fish are advantageous for comparisons of pallial growth mechanisms, as they exhibit an overall similar pallial morphology. We and others have extensively characterized neural progenitor lineages underlying pallial construction in zebrafish (Dong et al., 2012; Dirian et al., 2014; Furlan et al., 2017). In this species, the apical layer of cells, located along the T-shaped ventricle, contains all dividing neurogenic progenitors, while newly generated neurons sequentially stack in more basal layers (Furlan et al., 2017). In search for a comparative model with rapid pallial growth, we turned to the African turquoise killifish *Nothobranchius furzeri*. *N. furzeri* body size is similar to zebrafish at sexual maturity (~25mm), but it reaches this stage in 5 weeks, when it takes 3 months to zebrafish (Blažek et al., 2013). *N. furzeri* evolved this extremely fast post-hatching larval development as an adaptation to its ephemeral savannah pools habitat, to engender a new generation before ponds dry out (Blažek et al., 2013; Vrtílek et al., 2018). To validate our choice, we first set out to locate neural progenitors during the post-hatching and juvenile growth period. We used as markers Sox2, a highly conserved neural progenitor master transcription factor, and the proliferation marker Proliferating Cell Nuclear Antigen (PCNA) (Figure 1A). We observed that Sox2-positive and PCNA-positive cells localize at the apical surface, lining up the dorsally-opened pallial ventricle. They form a monolayer of cells, overlying differentiated neuronal cells located more basally in the parenchyme (Figure S1A). The growing killifish pallium therefore displays a centrifugal cellular organization, with a confinement of progenitors at the dorsal apical surface, as in zebrafish. However, as expected from the reported fast body growth, the killifish pallium grows very rapidly over the two weeks from hatching to 14 days post-hatching (dph). With linear increase, its surface reaches at 14 dph that of a 1.5 month-old juvenile zebrafish, hence growing 3 times faster (Figure S1B). While the pallial surface increases 5 times over these two weeks (Figure S1B), the cellular density of Sox2-positive cells remains rather constant, with only a 20 % decrease between 5 dph and 14 dph (Figure S1C). Thus, fast pallial growth in killifish is linked with an increased production of neural cells. Together, these observations indicate that the killifish is a suitable comparative model to explore adaptations that mediate accelerated neurogenesis and pallial growth.

**Figure 1:**
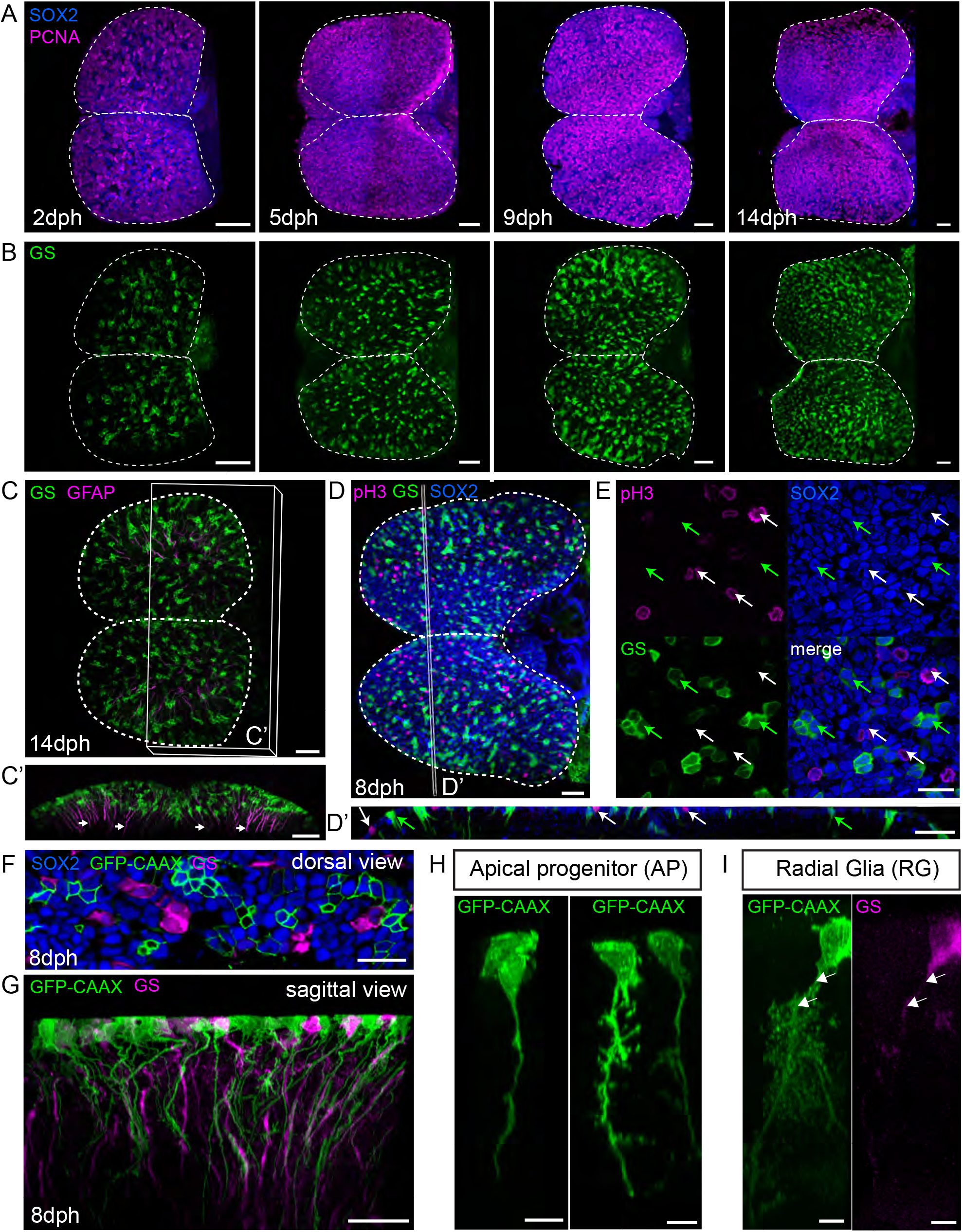
Two distinct progenitor types at the pallial surface of the killifish larval pallium. **A** Dorsal 3D-views of killifish pallium (anterior left) at 2, 5, 9 and 14 days post-hatching (dph) with a whole-mount immunostaining for Sox2 (blue) and PCNA (magenta) highlighting neural progenitors. A dotted line contours the two pallial hemispheres. **B**. Dorsal 3D-views of the same brains as in A, showing immunostaining signal for GS (green) to identify RG. **C-C’**. Immunostaining for GS (green) and GFAP (magenta) at 14 dph indicating GFAP-enriched process (arrows). **C**: dorsal 3D-view and **C’**: frontal view of the 3D reconstruction with a transverse hemisection along the plane shown in C **D-D’**. Immunostaining for GS (green), Sox2 (blue) and pH3 (magenta) at 8 dph. **D**: Dorsal 3D-view and **D’**: 5 µm transverse section through the 3D reconstruction **E**. High magnification of a single optical z-plane of the brain shown in D. White arrows in D’ and E point to non-glial AP progenitors entering mitosis at the apical surface and green arrows point to sparsely distributed RG. **F**. Optical z-plane showing AP cells electroporated with a GFP-CAAX-encoding plasmid (green), together with an immunostaining for GS (magenta) and Sox2 (blue). **G**. Sagittal view through a 3D reconstruction, highlighting the presence of a basal process on electroporated AP (green) and GS-positive RG (magenta). **H**. Examples of AP cells morphology. **I**. Typical RG morphology, showing the ramified pattern of the basal process. Individual cells were manually detoured on a 3D reconstruction to visualize their morphologies. Scale bars: 50 µm (A, B, C, D, D’, G); 20 µm (E, F); 10 µm (H, I). See also Figure S1.

### Radial glia are under-represented among neural progenitors of the killifish pallium

During embryonic development in zebrafish, as in other Vertebrates, ventricular progenitors convert to radial glia cells (RG), which will generate the large bulk of neurons and glial cells (Kriegstein and Alvarez-Buylla, 2009; Than-Trong and Bally-Cuif, 2015). This transition is marked by the induction of expression of a set of astroglial-specific genes, also shared by differentiated astrocytes in mammals. Astroglial markers notably include components of the synaptic glutamate recycling system (glutamine synthetase (GS) and glutamate transporters (Glast and Glt-1)), GFAP (glial fibrillary acid protein), BLBP (brain lipid binding protein) and S100b (calcium-binding protein) (Molofsky and Deneen, 2015). In the killifish pallium, immunohistochemistry for GS highlights the presence of RG among ventricular Sox2-positive cells at all stages analysed (Figure 1B). These cells display a radial morphology, with a long basal process enriched in GFAP protein (Figure 1C). They also express *slc1a2a* (Glt-1a), BLBP and S100b (Figure S1D-F). Surprisingly however RG do not cover the entire pallial surface and instead distribute sparsely in small groups (Figure 1D-E). This is in remarkable contrast with zebrafish, in which RG cells are the major constituents of the pallial ventricular surface and less than 10 % of Sox2-positive cells appear devoid of glial marker expression during post-embryonic stages (Figure S1G-I).

### Non-glial apical progenitors make the majority of the killifish pallial germinal zone

Killifish RG cells are intermingled with an additional pool of Sox2-positive progenitors (Figure 1D-E). Staining for pH3 (phospho-histone 3) indicate that the latter enter into mitosis at the apical surface (Figure 1D-E) and we will refer to them thereafter as apical progenitors (AP). To unravel the morphology of AP, we electroporated into the brain ventricle a plasmid encoding a membrane-bound GFP (GFP-CAAX). Viewing Sox2-positive AP from the dorsal (ventricular) side revealed that these cells possess an apical membrane domain (Figure 1F). This was confirmed by immunostaining against the tight-junction protein ZO-1 (zona occludens-1) (Figure S1J). Unexpectedly, we found that AP, like GS-positive RG, possess a long basal process (Figure 1G). Thus, AP show a radial morphology and an apico-basal polarity. However, we noticed that AP processes appear rather straight, with only few short branches (Figure 1G-H). In contrast, RG processes show a much more ramified pattern (Figure 1I), reminiscent of protoplasmic astrocytes, as previously noticed for zebrafish adult RG (Poskanzer and Molofsky, 2018). Thus, two types of neural progenitors compose the ventricular surface of the killifish pallium: prototypical RG, expressing a full set of astroglial markers, and radial AP, negative for astroglial markers, and with a more immature morphology.

### AP account for the majority of proliferating progenitors in the killifish pallium

Pallial growth in zebrafish is driven by the continued neurogenic capacity of RG (Dirian et al., 2014; Furlan et al., 2017), which present a sustained cycling activity during larval and juvenile stages (Figure S2A-B). Accelerated pallial growth in killifish may therefore result from intensified RG cycling activity. Alternatively, it may also involve the recruitment of additional neurogenic lineages. To test the first hypothesis, we analysed the proliferation rates of RG and AP in the killifish pallium along the first two weeks post-hatching, using the PCNA marker (Figure 2A). Larvae were also subjected to a 4-hour EdU pulse prior to fixation to label cells in S-phase. Our quantifications reveal that AP and RG differ significantly in their temporal proliferation profile (Figure 2B). The majority of AP are labelled by PCNA for the whole period; only a minor decrease of proliferation is observed at late time points. RG cells are already less proliferative than AP at 2 dph (54,1 % ± 4,1 % vs 97,3 % ± 0,6% of PCNA-positive RG vs AP, respectively, p<0.001). In addition, RG proliferation rate drops dramatically after 9 dph, reaching very low levels at 14 dph (14,9 % ± 3,4 %). Among PCNA-positive cells, ratios of EdU-labeled cells tend to decrease over time in both AP and RG without revealing any statistical differences between the two types of progenitors (Figure 2C). This suggests a parallel trend towards increasing length of growth phases of the cell cycle as pallial development progress. Altogether these data indicate that AP cycle actively and continuously during the post-hatching growth phase, while RG markedly reduce their proliferative activity.

**Figure 2:**
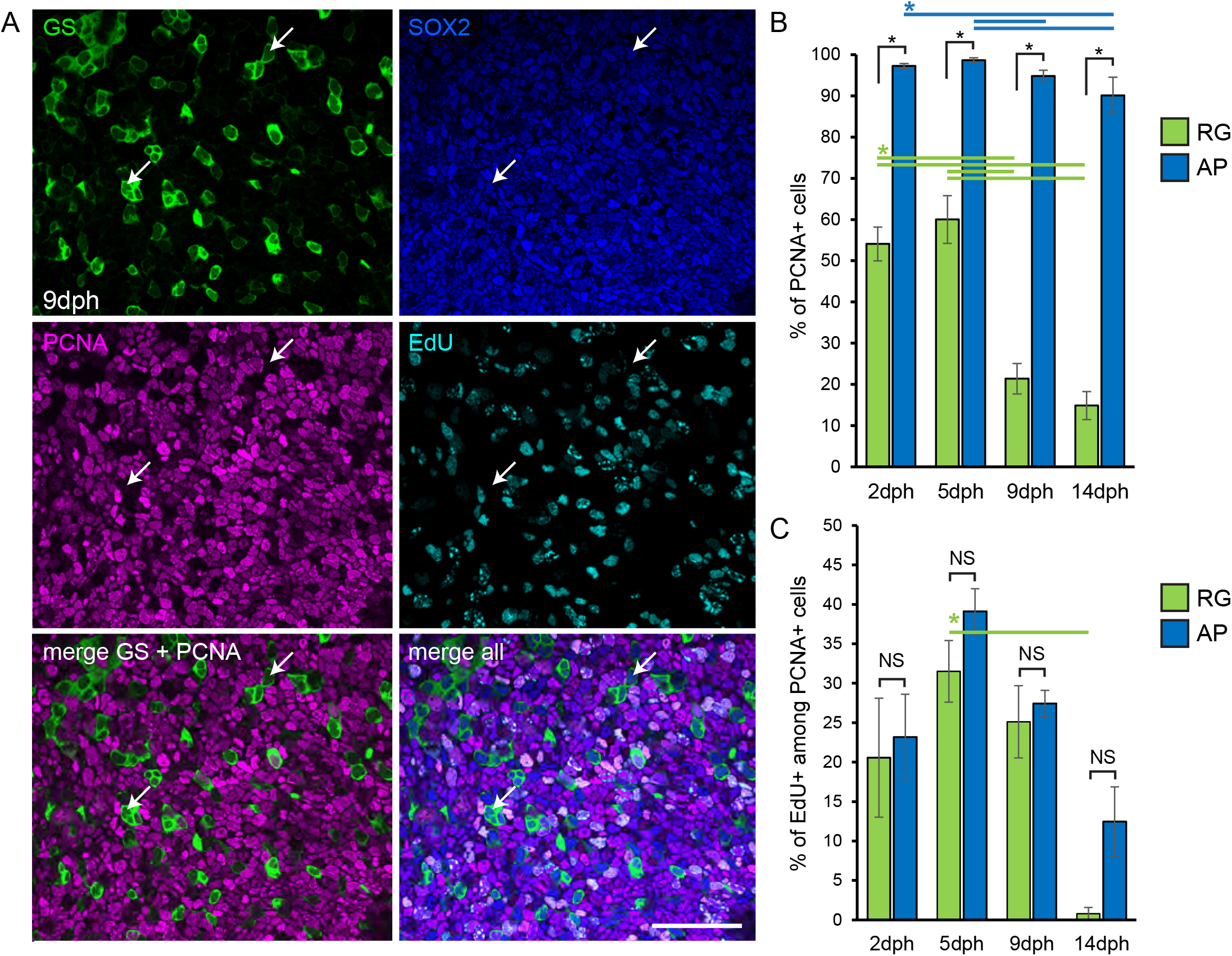
AP sustain a robust proliferative activity, while RG undergo a marked reduction of their activity. **A**. Optical z-plane showing immunostaining for GS (green), Sox2 (blue) and PCNA (magenta), and EdU detection (cyan), on the high magnification of the pallial surface at 9 dph. White arrows point to non-cycling RG cells. Scale bar: 50 µm. **B**. Proportion of PCNA-positive cells among RG (green bars) and AP (blue bars) at 2, 5, 9 and 14 dph. **C**. Proportion of EdU-positive cells among PCNA-positive cells among RG (green bars) and AP (blue bars). (*) corrected p-value<0.05; two-way ANOVA, followed by pairwise comparisons using Holm’s procedure. Data were rank transformed prior to analysis. Data are represented as mean ± SEM; n=6, 5, 5 and 3 for 2, 5, 9 and 14 dph respectively. See also Figure S2.

### RG enter premature Notch-dependent quiescence in killifish

Reduction of RG proliferation in killifish juveniles is reminiscent of the adult stage in the zebrafish pallium, when RG transition to a quiescent state, only occasionally activating to cycle (Chapouton et al., 2010). Alternatively, it could correspond to a definitive cell cycle exit for terminal glial differentiation. To decipher between these hypotheses, we first analysed the expression of highly conserved molecular players of the RG quiescence/activation cycle. Zebrafish adult RG cells are maintained in quiescence by the specific activation of one Notch receptor ortholog, Notch3 (Alunni et al., 2013; Chapouton et al., 2010; Than-Trong et al., 2018). A comparable function of Notch3-mediated signalling has been demonstrated in adult mouse neural stem cells (Ehret et al., 2015; Kawai et al., 2017). Upon exit of quiescence, in both mouse and zebrafish, activated cells induce the expression of the proneural gene *ascl1* (Andersen et al., 2014; Than-Trong et al., 2018). We therefore analysed the expression pattern of killifish orthologs of *notch3*, *ascl1* (*ascl1a* and *ascl1b*) and of the canonical Notch target *her4.2* (ortholog of *Hes5* in mammals). All these genes are expressed at the pallial ventricle in a salt-and-pepper fashion (Figure S3A). Using fluorescent ISH, we compared their transcripts distribution to the localization of GS-positive RG. We observed that *notch3* expression is confined to RG (Figure 3A). *her4.2* expression is highly enriched in RG, although a lower signal is detected in some AP. Finally, only few scattered RG express *ascl1* orthologs. These expression profiles are thus compatible with killifish RG being subjected to Notch3-dependent quiescence.

**Figure 3:**
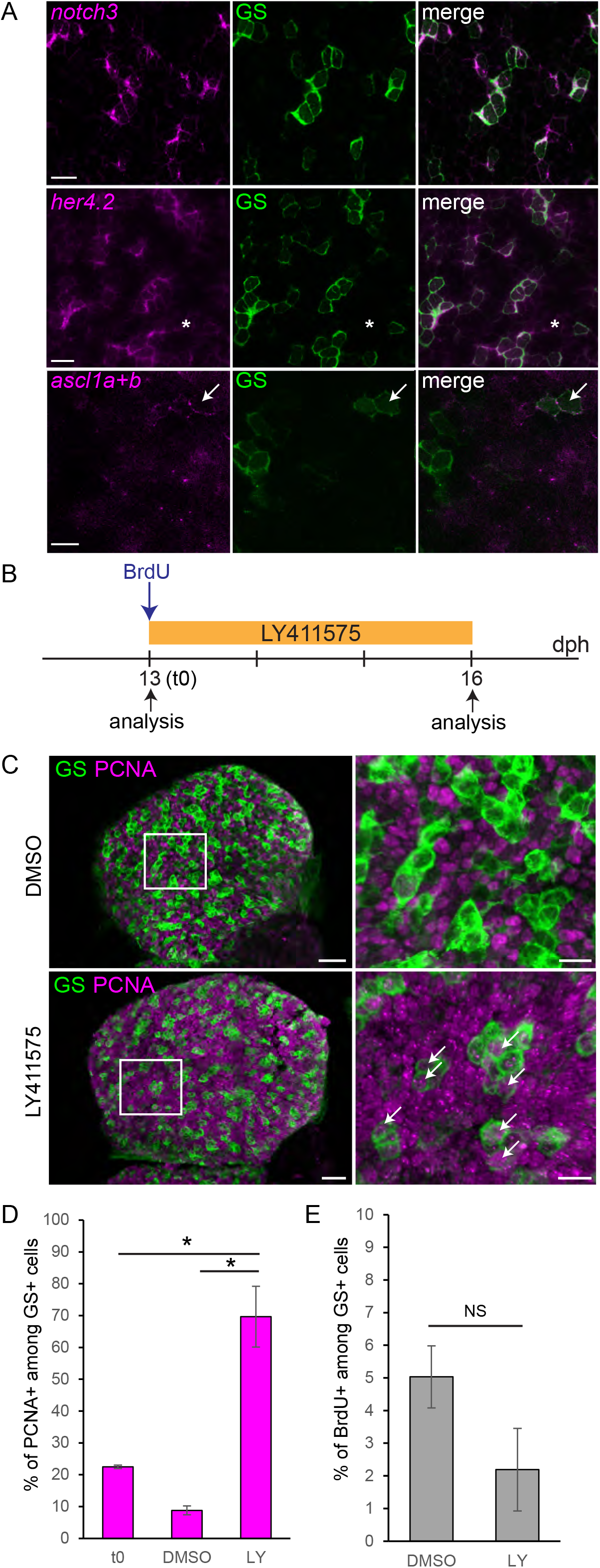
Killifish RG enter in a Notch3-dependent quiescent state. **A**. ISH for *notch3*, *her4.2* and *ascl1* orthologs (*ascl1a+b*) (magenta) combined with GS immunostaining (green) at 8 dph. Images are high magnifications of the pallial surface on a single optical z-plane. Asterisks indicate AP cells expressing low levels of *her4.2*. White arrows point to RG cells expressing *ascl1a/b*. Scale bar, 10 µm. **B**. Experimental design for the data shown in C-E. **C**. Double immunostaining for GS (green) and PCNA (magenta) in DMSO control (top panel) and LY-treated (bottom panel) brains, illustrating the increase in PCNA-positive RG cells (white arrows). Scale bars, 30 µm (left panels) and 10 µm (right panels). **D**. Proportion of PCNA-positive cells among GS-positive RG before treatment (t0) and after DMSO or LY treatment. **E**. Proportion of BrdU-labelled cells among RG after DMSO or LY treatment. (*) corrected p-value<0.05; one-way ANOVA, followed by pairwise comparisons using Holm’s procedure. Data were rank transformed prior to analysis. Data are represented as mean ± SEM; n=3 for each treatment condition. See also Figure S3.

In zebrafish, application of the gamma-secretase inhibitor LY411575 (LY) in the fish water efficiently blocks Notch activity and leads to a massive cell cycle re-entry of quiescent RG cells (Alunni et al., 2013). We thus tested whether LY would trigger a similar effect on killifish juvenile RG cells. We first verified that LY effectively inhibits the expression of the Notch target *her4.2* (Figure S3). Notably *her4.2* down-regulation is paralleled by an increased expression of *ascl1* orthologs, already suggesting an activation of RG cells. To further probe this, we treated 13 dph larvae for 3 days with LY (or DMSO as a control) and quantified the proportion of PCNA-positive RG cells (Figure 3B). In control brains, the proportion of PCNA-positive cells stays very low, in line with the low cycling activity of RG cells at this stage (Figure 1C, D). Under Notch blockade in contrast, the proportion of PCNA-positive cells drastically increases (DMSO: 8,8 ± 2,5 %; LY: 69,7 ± 9,6 % SEM, p=0,0027). In this experiment, we also applied a pulse of BrdU prior to treatment, followed by a chase (Figure 3B). In both control and LY treatment conditions, most RG cells are devoid of BrdU staining after the chase (Figure 3E). This demonstrates that activated RG upon LY treatment do not derive from previously cycling cells. Rather, they re-entered the cell cycle upon Notch blockade. We conclude that killifish RG cells enter a Notch-dependent quiescence state, similarly to adult zebrafish RG. However, they do so surprisingly precociously, in a context in which the pallium is still rapidly growing.

### AP account for the bulk of neurogenesis and pallial growth in killifish

Since RG enter into quiescence early, neuronal production in the killifish pallium likely relies mostly on their actively dividing neighbours, AP. We therefore examined AP fate potential and self-renewing capacity. We specifically traced their progeny by applying a pulse of EdU at 13 dph (Figure 4A). At this stage, because of their high proliferation rate relative to RG, AP indeed constitute the majority of EdU-incorporating cells (Figure S4). Because some AP express the Notch target *her4.2*, we analysed their fate both in control conditions and upon Notch blockade. Following the EdU pulse, fish were thus incubated for 3 days in DMSO or LY. After a second pulse with BrdU, fish were returned to fish water for an additional period of 3 days. In control conditions, we observed, after 3 days of chase, Sox2-negative EdU-labelled cells located basally in the parenchyme that we identify as neurons (Figure 4B-C). Noteworthy a substantial fraction of EdU-labelled cells is maintained as Sox2-positive AP after 3 and even 6 days of chase (68,2 ± 11,9 % and 31,9 ± 12,6 %; Figure 4C), and still incorporates BrdU after 3 days (Figure S4C). In contrast, the number of RG cells labelled by EdU is very low and constant over the whole period (Figure 4D). This tracing analysis overall shows that AP are both neurogenic and -at least over this experiment time frame - self-renewing. AP however do not generate RG, suggesting that they form an independent lineage.

**Figure 4:**
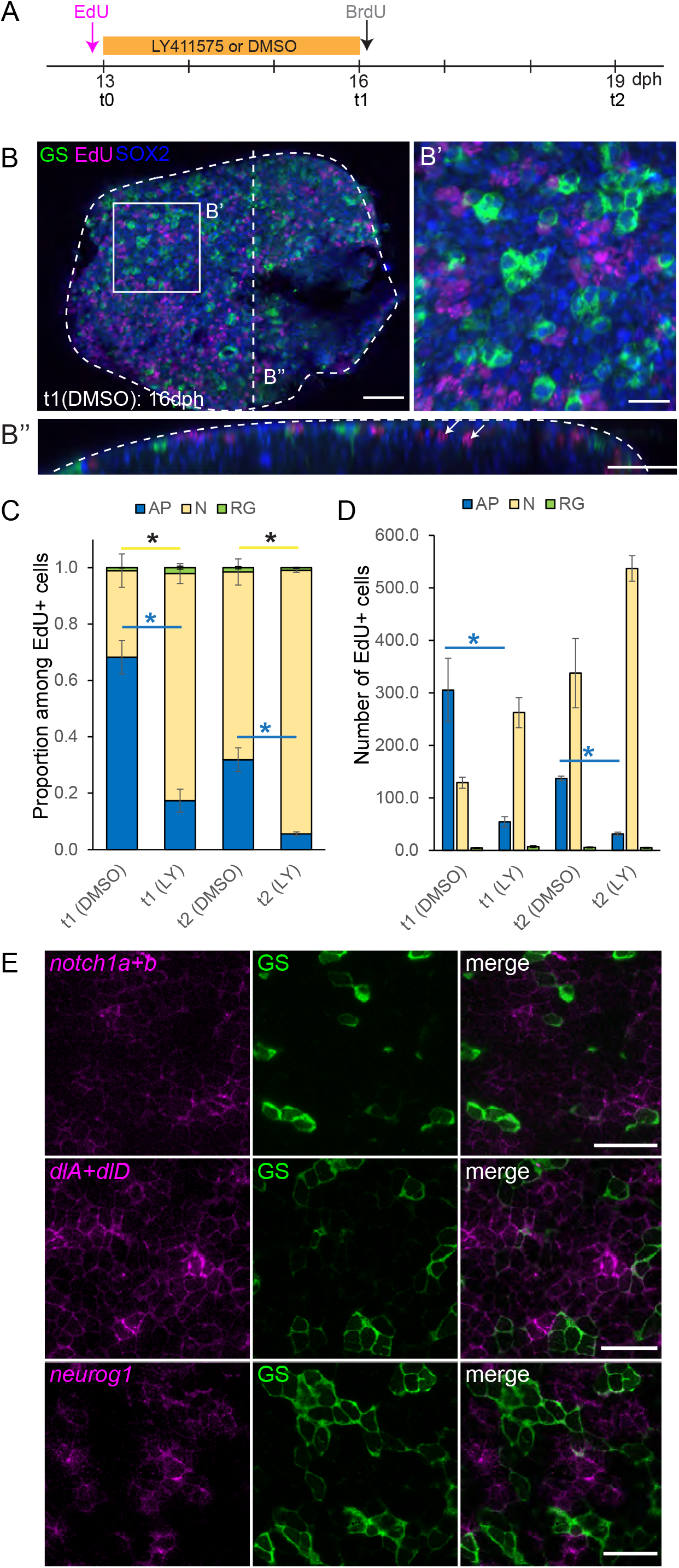
AP behave like embryonic progenitors. **A.** Experimental design. **B-B”**. Immunostaining for GS (green), Sox2 (blue) combined with EdU detection (magenta) on a DMSO control 16 dph killifish pallium. B’ is a high magnification of the dorsal view of the 3-D reconstruction shown in B. B” is 5 µm transverse section through the 3D reconstruction shown in B. White arrows in B” point to EdU-positive cells in the parenchyme that we identified as neurons. Scale bars, 50 µm (B, B”) and 20 µm (B’). **C**. Proportion of EdU-labelled cells with AP (blue, Sox2+GS-), neuronal (yellow, Sox2-GS-) or RG (green, Sox2+GS+) identity. (*) corrected p-value<0.05. Data were analysed for each cell type with a two-way ANOVA, followed by pairwise comparisons using Holm’s procedure. Proportions were arcsine transformed prior to analysis. Data are represented as mean ± SEM; n=2 (t1) and n=3 (t2) for each treatment condition for each treatment condition at t1 and t2 respectively. **D**. Number of EdU-labelled cells per counted area with AP (blue, Sox2+GS-), neuronal (yellow, Sox2-, GS-) or RG (green, Sox2+GS+) identity. (*) corrected p-value<0.05. Data were analysed for each cell type with a two-way ANOVA, followed by pairwise comparisons using Holm’s procedure. Data was rank transformed prior to analysis. Data are represented as mean ± SEM; n=2 (t1) and n=3(t2) for each treatment condition. **E**. ISH for *Notch1* orthologs (*notch1a+b)*, *Dll1* orthologs (*dlA*+*dlD*) and *neurog1* (magenta) combined with GS immunostaining (green) at 8 dph. Images are high magnifications of the pallial surface on a single optical z-plane. Scale bar, 20 µm. See also Figure S4.

### AP use the proneural gene *neurog1* and rely on Notch1 signalling for their maintenance, similarly to embryonic neural progenitors

In vertebrate neurogenic progenitors, Notch-mediated intercellular communication balances self-renewal and differentiation, ensuring their long-term maintenance. This function is mainly relayed by Notch1 receptors (Chambers et al., 2001; Hatakeyama and Kageyama, 2006; Basak et al., 2012; Alunni et al., 2013). The balance relies on antiphase oscillations in neighbouring progenitors of Hes-related genes on the one hand versus Notch-ligands and proneural genes on the other hand (Kageyama et al., 2008). In line with this, our fate analysis of AP reveals that the relative proportions of neurons and AP are disturbed by Notch inhibition (Figure 4C). This is mainly due to a significant decrease in the number of EdU-positive cells maintained as AP, paralleled by an increase in the production of EdU-positive neurons (Figure 4D). Consistently, we also show that AP preferentially express *notch1* orthologs (*notch1a* and *notch1b*). Additionally, we observed among AP a salt-and-pepper pattern of expression *her4.2* (Figure 3A), of Dll1-related Notch ligands (*dla* and *dld*) and of the proneural gene *neurog1* (Figure 4E and S4D). Of note, the expression of *neurog1* in AP contrasts with RG, which use *ascl1* orthologs as proneural factors when activated (Figure S3B), but also with zebrafish, in which *neurog1* expression is restricted to early embryonic stages. While its expression in the early embryonic zebrafish pallium has been previously reported (Blader et al., 1997; Korzh et al., 1998), we indeed did not detect any at post-embryonic stages (Figure S4E). Our results therefore indicate AP show characteristics of early-embryonic neurogenic progenitors that would be retained at larval stages.

## Discussion

Altogether our study unravels a striking mosaic heterochrony within the killifish pallial germinal zone, where progenitors at different stages of maturation cohabit: adult-like quiescent RG and embryonic-like neurogenic AP, the latter defined by their immature morphology and their reliance on the *notch1/neurog1* interplay for the maintenance of their progenitor properties. This contrasts with zebrafish, in which the pallial surface is composed mainly of RG acting as neural progenitors from embryonic to adult stages. The zebrafish situation corresponds likely to the ancestral condition in teleosts. Indeed we observed GS-positive RG cells covering the whole pallial surface in medaka fish (Figure S4F), a species more closely-related to killifish (75 my divergence (Betancur-R et al., 2013)). We propose, therefore, that the presence of an AP lineage is a killifish adaptation accommodating the explosive growth imposed by its ephemeral habitat.

We show that both AP and RG have an apical contact and a basal process; it raises the question of how these two types of progenitors can achieve such different temporal behaviours while residing in the same local niche. This must rely on a differential sensitivity or accessibility to temporal cues, but the underlying mechanisms remain to be assessed. Our results also indicate that in killifish, in contrast to other Vertebrates, only a fraction of pallial progenitors switch on the astroglial program. What licences this mosaic induction remains to be further investigated. But since the dual composition of the pallial germinal zone is already present at the pre-hatching diapause III stage (not shown), e.g. prior to the rapid growth phase, it is not a mere consequence of an intensification of growth-promoting signals. Rather, an early change in the encoded neural progenitors program must have occurred in the killifish lineage.

The killifish is a newly co-opted vertebrate model for aging studies, due to its exceptionally short lifespan (Cellerino et al., 2016; Harel and Brunet, 2015). Recent studies are beginning to explore causal factors of its accelerated aging, including in the brain (Harel et al., 2015; Cui et al., 2019; Tozzini et al., 2012; Ripa et al., 2017; Matsui et al., 2019). The fast ageing in *N.furzeri* is hypothesized by some authors to be a trade-off to rapid larval and juvenile growth, but this lead remains mechanistically unexplored (Reichard and Polačik, 2019). Killifish are not paedomorphic and are considered to simply globally speed up body development (Reichard et al., 2015). However, we demonstrate here that developmental timing at the individual organ level is actually impacted. This may have profound implications for pallial structure and physiology. Indeed, in both vertebrate and invertebrate models, neural progenitors have been shown to sequentially produce distinct neuronal subtypes during development (Rossi et al., 2017). The maintenance of embryonic-like AP progenitors in killifish could therefore have major consequences on the neuronal cell types composing the pallium. The sparse distribution of RG at the pallial surface also implies a reduced astroglial to neurons ratio. Considering the prominent role of astroglial cells in neuronal function, this change could deeply impact brain physiology (Barres, 2008) but also contribute to brain ageing phenotypes (Palmer and Ousman, 2018).

Interspecies alterations in neural progenitor lineages have been observed in mammals, although to control brain size. In mammals, the expansion of the pallium during evolution is thus driven by the emergence and diversification of basally-located progenitors, which amplify neuronal production downstream of apical RG (Romero and Borrell, 2015). While no basal progenitors are present in the pallium of birds, reptiles and teleost fish, some have been observed in a shark, highlighting a flexibility in pallial lineage patterns (Docampo-Seara et al., 2018). The strategy to amplify basal progenitors to boost neuronal production in mammals differs from what we observed in killifish. However, it shares a common feature. Indeed, in both cases, neuronal production is augmented without altering RG division rate. This suggests an inherent constraint imposed to vertebrate brain development, demanding the maintenance of an unwavering RG population. In conclusion our work reveals substantial modifications of anterior brain development associated to increased developmental speed, and highlights nodal points of evolutionary plasticity and constraints in vertebrate neural progenitor lineages.

## Acknowledgements

We thank members of the L. B-C. lab for their critical input, S. Bedu for providing expert zebrafish care and Kasia Banasiak for help in data collection. We thank Sylvie Rétaux for her careful and critical reading of the manuscript. We thank Alessandro Cellerino for sharing his expertise in the killifish model. We are grateful to Sylvie Schneider-Maunoury (IBPS) for her support in the establishment of the killifish facility. We thank Matthieu Simion (NeuroPSI) for providing Medaka larval brains. M.C. is supported by INSERM. Work in the L. B-C. lab was funded by the ANR (grant ANR-2012-BSV4-0004-01), Labex Revive, Centre National de la Recherche Scientifique, Ecole des Neurosciences de Paris (ENP), Institut Pasteur and the European Research Council (AdG 322936).

## Author Contributions

Conceptualization, M.C. and L.B.-C.; Investigation: M.C., M.L.; Resources, A.M., B.H. and M. B.; Supervision, M.C. and L.B.-C.; Writing - Original draft, M.C.; Writing – Review and Editing, M.C. and L.B.-C.; Funding Acquisition, L.B.-C.

## Declaration of Interests

The authors declare no competing interests.

## STAR Methods

### Key Resources Table

KEY RESOURCES TABLE

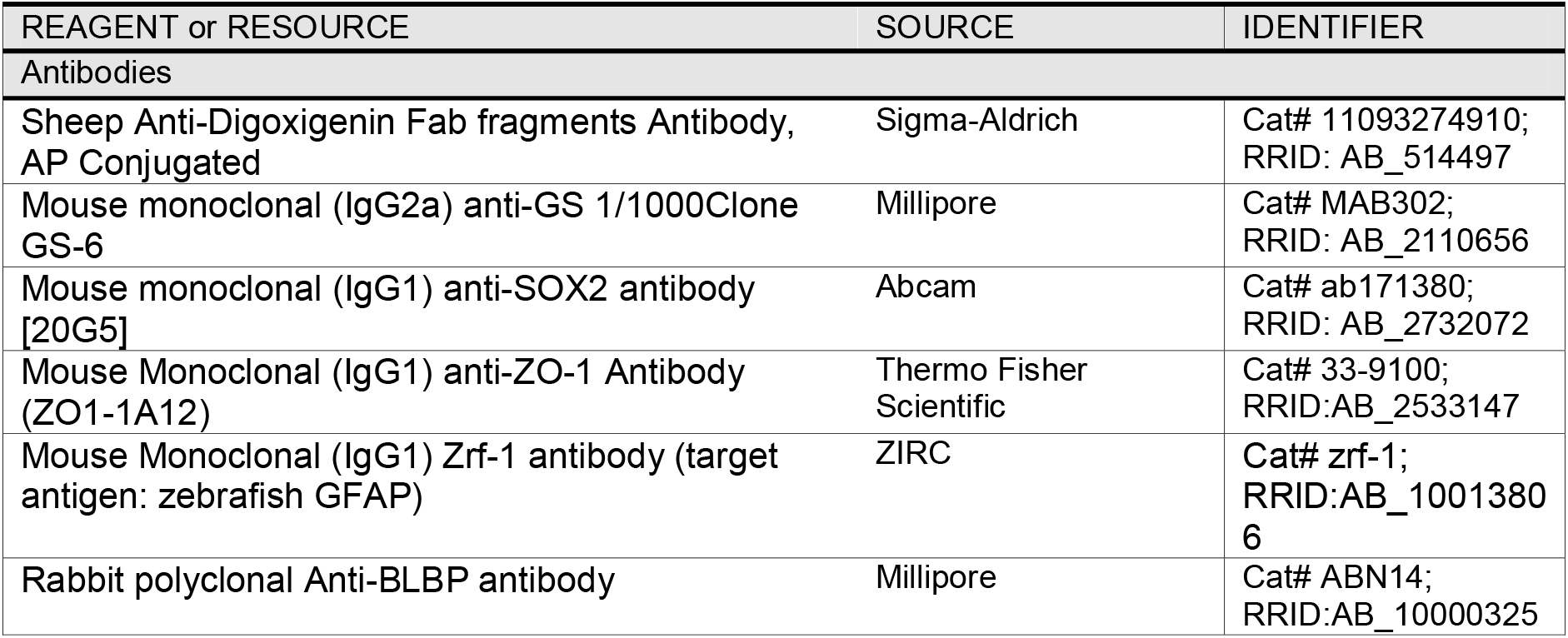
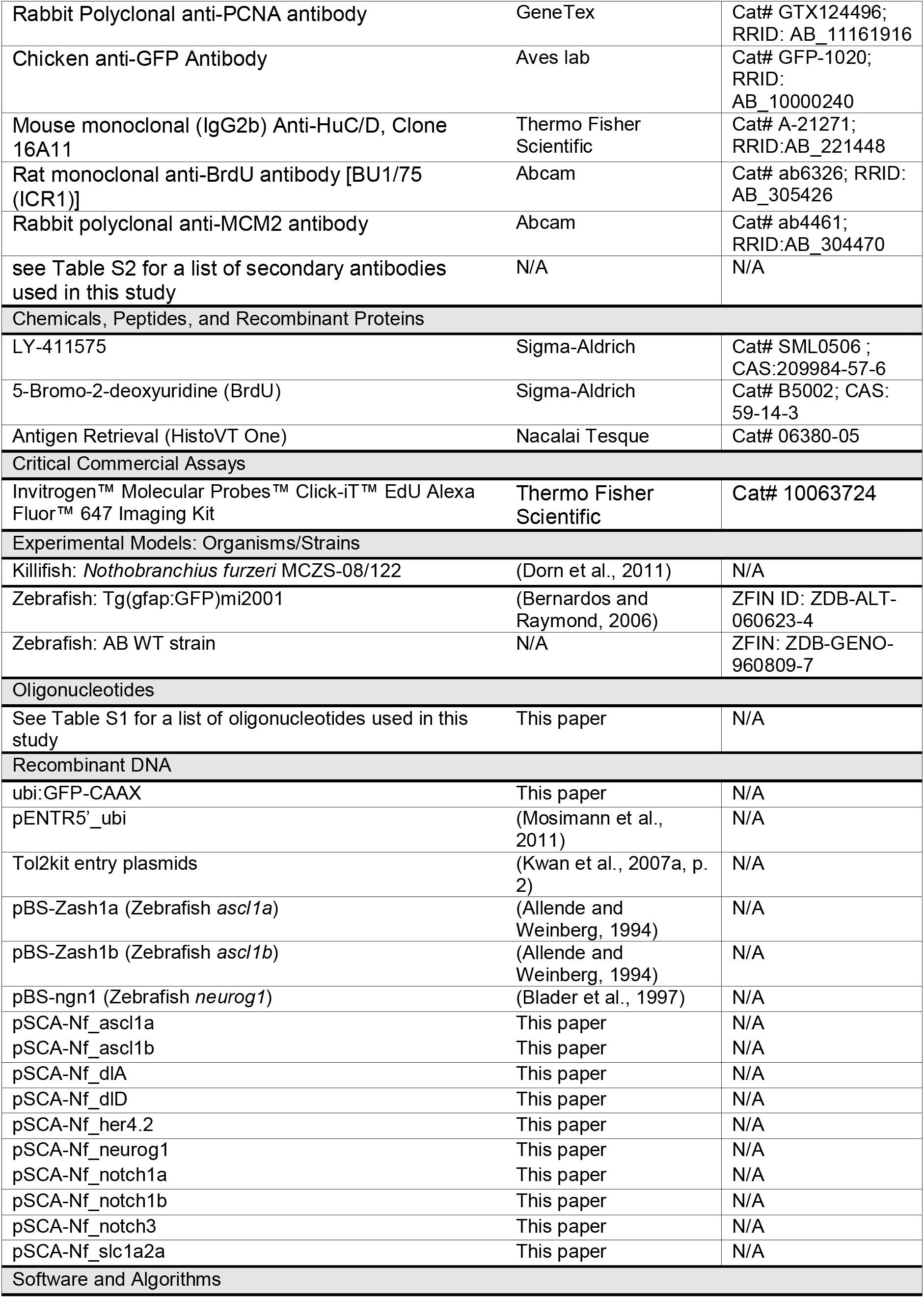
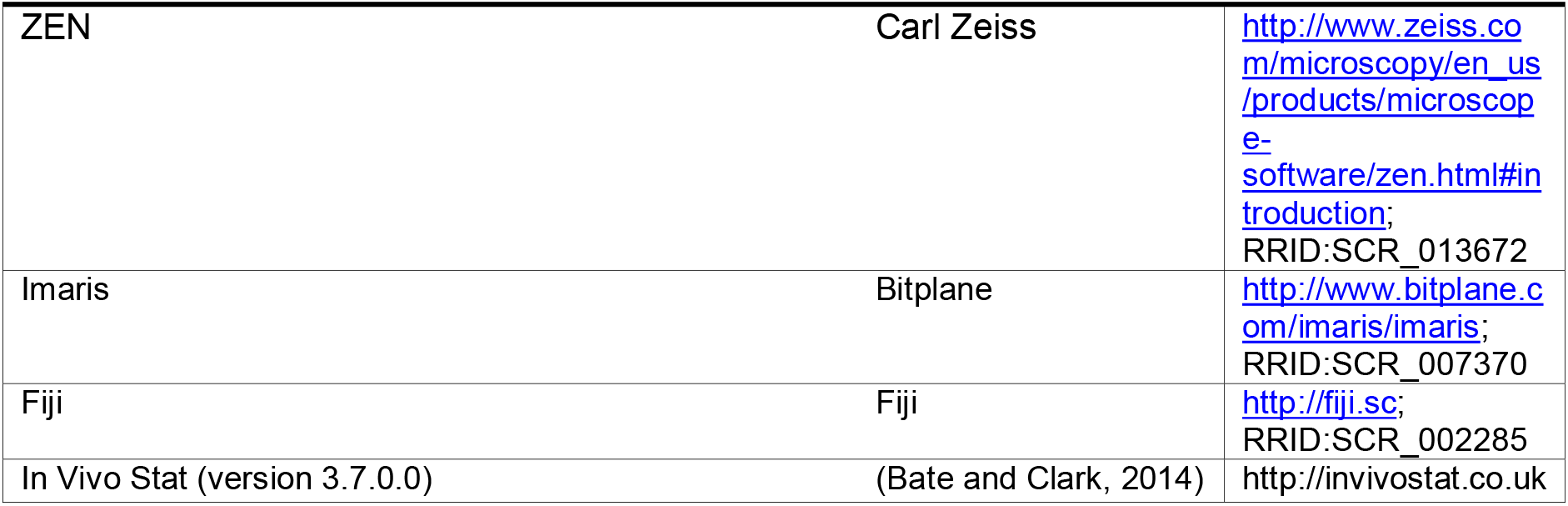

### Contact for Reagent and Resource Sharing

Further information and requests for resources and reagents should be directed to and will be fulfilled by the Lead Contact, Laure Bally-Cuif.

### Experimental Model and Subject Details

#### Killifish

The *Nothobranchius furzeri* strain MZCS‐08/122 was used for experiments (Dorn et al., 2011). Adult fish were fed with frozen red bloodworms (*Chironomidae*) once a day. Fish were kept in 8L breeding tanks housing one male and two females. Breeding tanks were equipped with sand boxes from which eggs were collected once a week. Eggs were bleached using a solution of Sodium hypochlorite (78µL of NaOCl (Sigma, 425044) diluted in 100 mL of 0.3x Danieau’s Medium). Embryos were placed on petri dishes filled with moist coconut fibre and stored in the dark at room temperature. In these conditions, embryos arrest in diapause II. Plates were checked at least twice a week to remove dead embryos and avoid fungal contamination. To stimulate diapause II exit, plates were placed in an incubator at 26 °C with a 12h/12h light/dark cycle until the embryos reached the diapause III stage, characterized by the “golden eyes” phenotype (Polacik et al., 2016). For hatching, a peat moss extract solution was prepared: boiling water was poured on coconut fiber, and the solution was filtered on filter paper and autoclaved. Eggs were placed in a small plastic tray and immersed in cold peat moss extract. To ensure sufficient oxygenation, oxygen tablets were broken into small pieces and added to the hatching tray. Hatched larvae were transferred to a larger box with system water and immediately fed with freshly-hatched brine shrimp (*Artemia*) twice a day. Larvae were raised at 26 °C, with a daily 50 % water change, and fed twice a day with artemia. Water levels were progressively increased as fish were growing.

#### Zebrafish

Wild-type zebrafish from the AB line were used for the experiments. Zebrafish were maintained using standard fish-keeping protocols and staged according to (Parichy et al., 2009).

#### Medaka

Fixed 10 dpf medaka larvae were kindly provided by Matthieu Simion (J.S. Joly laboratory, NeuroPSI, Gif-sur-Yvette).

All animal experiments were carried out in accordance with the official regulatory standards of the Department of Paris (agreement number B75-15-22 to L.B.-C. and A-75-05-25 to A.M.) and conformed to French and European ethical and animal welfare directives (project authorization from the Ministère de l’Enseignement Supérieur, de la Recherche et de l’Innovation to L.B.-C.).

### Method Details

#### Tissue fixation

Fish larvae were euthanized with tricaine on ice and fixed overnight at 4 °C in a 4 % solution of paraformaldehyde (PFA) in PBS. After two PBS washes, brains were dissected out in PBS and bleached with a peroxide-containing bleaching solution (5 % Formamide, 0.5X SSC, 3 % H_2_0_2_) until dark pigments are no longer visible. Brains were subsequently dehydrated in serial dilutions of methanol/PBS. Brains were stored in 100 % methanol at −20 °C until use.

#### BrdU and EdU pulse labeling

BrdU and EdU were applied in the fish swimming water for 4h, except for the experiment presented in figure 4 were the EdU was applied for 24h. The final concentrations used were 1 mM for BrdU and 400 µM for EdU. After the pulse, fish were transferred to a tank with fresh fish water during chase periods.

#### LY411575 treatment

Stock solutions of LY411575 (LY) at 10 mM were prepared by solving 5 mg of LY in 1.05 mL DMSO and stored at −80 °C until use. To block Notch signalling, LY was applied in the fish swimming water at a final concentration of 10 µM. The solution was refreshed every 24 h. Control fish were treated with the same final concentration (0.1 %) of DMSO carrier.

#### Electroporation

Fish larvae were anesthetized with tricaine in system water. With a thin needle, a solution containing the DNA plasmid construct (concentration 2 µg/µL) was injected into the forebrain ventricle. The DNA construct was a vector encoding a membrane targeted GFP (GFP-CAAX) under the control of the zebrafish *ubiquitin* promoter (Mosimann et al., 2011). This construct was generated by performing a gateway LR reaction (LR II clonase, Invitrogen) using the pENTR5’_*ubi* (Mosimann et al., 2011) pME-GFP-CAAX, p3E-polyA and pDest-Tol2pA2 entry vectors (Tol2kit (Kwan et al., 2007b, p. 2)). After intraventricular injection of DNA, three short electropulses (width 50 ms, space 500 ms) at 50 mV were immediately applied. Fish were transferred to system water for recovery and sacrificed 24 h later for immunohistochemistry.

#### In silico searches and primer design for cloning

For each candidate gene, the protein sequence from the zebrafish ortholog was retrieved from the NCBI and blasted against transcriptomic database of *Nothobranchius furzeri* (http://nfintb.leibniz-fli.de/nfintb/blast.php). Potential orthologs were selected based on E-value and reverse blast hits results. Orthology relationships were confirmed by phylogenetic analyses using available Vertebrate protein sequences and a Maximum-Likehood tree reconstruction method (not shown) (http://www.phylogeny.fr/). Primers were designed using the primer3 software (http://biotools.umassmed.edu/bioapps/primer3_www.cgi).

#### RNA extraction

15 dph killifish brains were dissected in cold PBS and placed in an Eppendorf tube. Brain tissue was homogenized with a pestle in Trizol solution (Thermofisher 12044977, 1 mL Trizol for 50 – 100 mg tissue). After a five-minute incubation at RT, chloroform was added (200µL per mL of Trizol). The Eppendorf tube was shaken for 15 seconds and centrifuged at 12000 g for 15 min. The upper phase was transferred to a fresh tube containing 500 µL Isopropanol and incubated at RT for 10 min. The tube was centrifuged at 12000 g at 4 °C for 10 min. After removal of supernatant, the pellet was washed with 70 % EtOH and dissolved in 44 µL RNAse-free water (Ambion). Genomic DNA was removed using a RQ1 RNAse-free DNAse treatment (Promega): 5 µL of 10x DNase buffer and 1 µL of RNase-free DNase was added to the 44 µL of RNA and the mix was incubated for 30 min at 37°C. To stop the reaction, 1 µL of 0.25 mM EDTA was added and the tube was incubated at 65 °C for 10 min. RNA concentration was measured using a nanodrop and stored at −80 °C.

#### Reverse Transcription

DNAse-treated RNA was reverse transcribed with Superscript II reverse transcriptase (Invitrogen) using random hexamer primers and according to the manufacturer’s instruction. Briefly 1 µg of DNAse-treated RNA was diluted in 9 µL of RNAse-free water. 1 µL of hexamer random primers (100µM stock) was added and the mix was incubated at 70 °C for 5 min. Following the addition on ice of 4 µL of 5x buffer, 2 µL of DTT (0.1 M), 2 µL of dNTPs (10 mM each) and 1 µL of RNase inhibitor (Promega), the mix was incubated at 25 °C for 5 min. 1 µL Superscript II enzyme was added, and reverse transcription was performed by incubation at 25 °C for 10 min, at 42 °C for 1h, and at 70 °C for 10 min. The cDNA was stored at −20 °C and used as a template for PCR reactions.

#### Cloning

*Nothobranchius furzeri* transcripts were amplified by PCR using primers listed in table S1 and either GoTaq (Promega) or Platinum Taq polymerase (Invitrogen). Fragments were cloned using Strataclone PCR cloning kit (Agilent technologies) according to the manufacturer’s instructions. Clones sequences were verified by sequencing with universal M13 primers (GATC biotech).

#### RNA probes synthesis

For RNA probes synthesis, 10 µg of plasmid DNA was linearized using the appropriate enzyme, purified using the NucleoSpin Gel and PCR Clean-up kit (Machery-Nagel) and eluted in 20 µL of RNAse-free water (Ambion). To generate DIG-labeled probes, 1 µg linearized plasmid was used as a template for transcription reaction using T3 or T7 RNA polymerase (Promega) and DIG-labeling mix (Roche) according to the manufacturer’s instructions. Transcription reaction was carried out for 3 h at 37 °C and a final incubation with Turbo DNase I was performed to remove template DNA. Unincorporated nucleotides were removed with the ProbeQuant G-50 Micro Columns Kit (GE Healthcare). The RNA probe was stored at −80 °C.

#### Whole-mount in situ hybridization

*In situ* hybridization were carried out according to standard protocols as previously described (Furlan et al., 2017). Zebrafish probes used in this study are *neurog1* (Blader et al., 1997), *ascl1a* and *ascl1b* (Allende and Weinberg, 1994). Killifish probes were generated in this study as described above (table S1). Briefly brains were rehydrated, washed with PBST and incubated in hybridization buffer at 65 °C for 3 h. Brains were then incubated overnight at 65 °C with DIG-labeled RNA probes diluted at 1/100 in hybridization buffer. Brains were washed in serial dilutions of hybridization buffer/2XSSC at 65 °C to remove excess probe. An immunohistochemistry with anti-DIG antibody coupled to alkaline phosphatase (Roche, 1/5000 dilution) was then performed. *In situ* signals were revealed either with NBT/BCIP (Roche) or Fast Red (Sigma) for fluorescent visualization.

#### Whole-mount IHC on brains

Brains stored in 100 % MeOH were rehydrated and washed 3 times with PBST (0.1 % Tween-20 in PBS). When immunostaining included a nuclear marker, an incubation in HistoVT One (Nacalai Tesque) buffer for 1 h at 65 °C was performed. If the immunostaining included BrdU staining, an incubation in HCl 2N at RT for 25 min was performed. Brains were washed three times for 5 min each with PBST. The brains were incubated into Blocking Solution (5 % Normal Goat Serum, 0.1 % Triton X-100 in PBS) for at least 1 h at RT. Primary antibodies were diluted in blocking Solution and incubated for 24 h at 4 °C. Brains were subsequently washed six times for 15 min with PBST and incubated overnight at 4 °C with secondary antibodies diluted 1:1000 in Blocking Solution (see Table S2). The brains were then washed six times for 15 min in PBST. For EdU detection the Invitrogen™ Molecular Probes™ Click-iT™ EdU Alexa Fluor™ 647 Imaging Kit was used, following the manufacturer’s instructions. Brains were mounted in Vectashield solution for confocal imaging.

#### Image acquisition

Fluorescent images of whole-mount telencephali were acquired on confocal microscopes (LSM700 and LSM710, Zeiss), using either a 20X air objective (Plan Apochromat 20x/0.8 M27) or a 40X oil objective (Plan-Apochromat 40x/1.3 Oil M27). For large samples, a tile scan was performed and stitching was done with the ZEN software after imaging. 3D renderings were generated using the Imaris software (versions 8 and 9, Bitplane). The 3D image was cropped to feature only the pallium as our region of interest and contrast was adjusted for each channel. For Sox2 immunostaining signals, background was substracted using the remove outliers function of the Fiji image analysis software. For *in situ* hybridization experiments revealed in NBT/BCIP images were acquired using a macroscope (Zeiss).

### Quantification and Statistical Analyses

3D Images were segmented manually using semi-automatic detection with the Imaris spots function followed by manual curation. For quantification of proliferation in zebrafish RG at post-embryonic stages, the whole pallium was segmented (Figure S2). For quantification of proliferating cells along killifish post-hatching development (Figure 2), one hemisphere per brain was segmented. For the other experiments counting was performed on a region of interest of the ventricular zone (69 × 122 µm and 170 × 240 µm for figure 3 and 4 respectively). Data are presented as mean ± standard error of the mean (s.e.m.). Statistical analyses were carried out using InVivoStat (Bate and Clark, 2014; Clark et al., 2012) and graphical plots were generated using Microsoft Excel. The normality of responses was assessed using normality probability plots and the homogeneity of the variance was inspected on a predicted versus residual plot. When the responses deviated noticeably from either criterion, they were arcsine or rank-transformed. Data displaying an approximately Gaussian distribution of residuals and homoscedastic responses with or without transformation were analysed using parametric tests. Overall factor effects were determined by analysis of variance (ANOVA) and pairwise comparisons were carried out with least significant difference tests (LSD). P-values were adjusted for multiple comparisons according to the Holm’s procedure. All the statistical tests performed were two-tailed and their significance level was set at 5 % (α=0.05).

## Supplemental Information

**Figure S1: Two types of neural progenitors reside at the apical surface of the rapidly growing killifish pallium (related to figure 1)**

**A.** Immunostaining of a 7dph killifish pallium for the markers Sox2 (blue), PCNA (magenta) and HuC/D (green). HuC/D (encoded by *elavl3/4* genes) is a marker of differentiated neurons. Dividing neural progenitors are restricted to the ventricular surface. The top panel shows a dorsal view of a 3D reconstruction and bottom panels correspond to a 5µm transverse section through the 3D. Scale bar, 20 µm. **B**. Comparison of pallial growth between the killifish *N. furzeri* and zebrafish. Pallial surface was estimated by measuring pallial maximal length (L, along the antero-posterior axis) and width (l, along the left-right axis). Measurements were performed on samples from pre-hatching diapause III stage (t0) to sexually adult stage (35d) in *N. furzeri* and from Fle stage (~5d post-fertilization, t0) to juvenile (~45d post-fertilization) stage in zebrafish. Zebrafish larvae were staged according to (Parichy et al., 2009) and plotted using the corresponding average developmental time in AB fish. (Fle: early flexion; CR: caudal fin ray appearance; AC: anal fin condensation; DC: dorsal fin condensation; aSB: anterior swim bladder appearance; DR: dorsal fin ray appearance; PB: pelvic bud appearance; PR: pelvin fin ray; SP: squamation onset posterior; J: Juvenile). **C**. Density of Sox2-positive cells at the pallial surface of the killifish. To estimate density, a surface was created on one pallial hemisphere using the Imaris surface function and the total number of Sox2-positive cells was quantified. (*) corrected p-value. Data were analysed with a one-way ANOVA, followed by pairwise comparisons using Holm’s procedure. Proportions were rank transformed prior to analysis. Data are represented as mean ± SEM; n=6, 5, 5 and 3 for 2, 5, 9 and 14dph respectively. **D**. ISH for *slc1a2a* (encoding an ortholog of glutamate transporter GLT-1, magenta) combined with GS immunostaining at 9dph. Images are dorsal views of a 3D reconstruction. Scale bar, 50µm **E**. Triple immunostaining for BLBP, Sox2 and GS at 9dph. Images show high-magnifications of the pallial surface on a single optical z-plane. Scale bar, 20µm **F**. Triple immunostaining for S100b, Sox2 and GS at 9dph. Images show high-magnifications of the pallial surface on a single optical z-plane. **G**. Proportion of non-glial Sox2-positive cells at the ventricular surface of the zebrafish pallium at the Juvenile (J) stage. Sox2-positive cells were quantified in the transgenic line Tg(*gfap:GFP*)^mi2001^, and cells without GFP signal were scored as non-glial progenitors. Data are represented as mean ± SEM; n=2. **H.** Proportion of AP at the ventricular surface of the killifish pallium at 14dph. Quantifications were performed on brains immunostained for GS, Sox2 and PCNA (see Figure 1A-B) and the Sox2-positive GS-negative cells are scored as AP. Data are represented as mean ±SEM; n=3 **I.** Direct comparison of the distribution of RG cells at the surface of the pallium in zebrafish at the juvenile stage (left panel) and killifish at 14dph (right panels) at the same magnification, using a double immunostaining for GS (green) and Sox2 (blue). A dorsal view of a 3D reconstruction is shown. Scale bar, 100µm for the global view and 20µm for high magnification views. **J**. Triple immunostaining for ZO-1 (gray), GS (green) and PCNA (magenta) at 14dph. Images are high-magnifications of the pallial surface from a 3D reconstruction. Scale bar, 20µm.

**Figure S2: In contrast to killifish, RG in zebrafish sustain proliferation during post-embryonic stages (related to Figure 2)**

**A**. Dorsal views of 3-D reconstruction of the zebrafish pallium at stage CR (~10dpf). Neural progenitors are labelled with Sox2 antibody (blue), RG are labelled by a GFP reporter of the Tg(*gfap*:GFP)^mi2001^ line and proliferating cells are labelled with a MCM2 antibody (magenta). Scale bar, 70µm. **B.** Proportion of MCM2-positive proliferating cells among the RG population (GFP-positive in the Tg(*gfap*:GFP)^mi2001^ line). RG cells maintain a quite stable proliferation rate across stages. Data are represented as mean ± SEM. The number of samples used for each stage is indicated on the graph. Comparisons between these stages show no statistically significant differences. Data were analysed with a one-way ANOVA, followed by pairwise comparisons using Holm’s procedure. Proportions were rank transformed prior to analysis.

**Figure S3: Expression of Notch pathway genes involved in the quiescence/activation cycle of RG (related to figure 3)**

**A**. Mosaic expression of *notch3*, *her4.2* and *ascl1* orthologs in the killifish pallium at 8dph. Whole-mount ISH of the pallium are shown as well as higher magnification in the top right corner. Some signal-positive cells are indicated by white arrows. A 50µm-vibratome section is also shown for *notch3*, showing that the expression restricted to the apical ventricular layer. Scale bar, 100µm **B**. Effect of Notch inhibition on the expression of *her4.2* and *ascl1* orthologs. Whole-mount ISH of the pallium are shown as well as higher magnification in the top right corner. *her4.2* signal totally disappears after 24h of LY treatment. Barely any signal for *ascl1a+b* is detected in control brains at 14dph, in keeping with the low activation rate of RG at this stage. In contrast, mosaic expression of *ascl1a+b* in some cells of the ventricular surface is detected after LY treatment (white arrows). Scale bar, 200µm

**Figure S4: Self-renewing and neurogenic AP maintain embryonic-like marker expression (related to figure 4)**

**A-A’.** Dorsal view of a killifish pallium at 13dph with an immunostaining for Sox2, GS and detection of EdU. A’ is a higher magnification of the boxed area in A. Most EdU-labeled cells are AP. Scale bar, 50µm **B**. Proportion of EdU-labeled cells that are AP (blue, Sox2+GS-) or RG (green, Sox2+GS+) immediately following the EdU pulse (t0). Data are represented as mean ± SEM; n=2. **C**. Proportion of EdU-labeled AP cells that have incorporated BrdU immediately following the BrdU pulse (t1) or after 3 days of chase. Data are represented as mean ± SEM; n=2 (t1) or n=3(t2) for each treatment condition. Statistical analysis does not reveal any statistical difference (2-way ANOVA approach followed by pairwise comparisons, data were ranked transformed prior to analysis). **D**. *notch1* orthologs, *Dll1* orthologs and *neurog1* expression in the killifish pallium at 8dph. Whole-mount ISH of the pallium are shown as well as higher magnification in the top right corner. Some signal-positive cells are indicated by white arrows. A 50µm-vibratome section is also shown for *neurog1*, highlighting the salt-and-pepper expression in the ventricular layer. Scale bar, 100µm. **E**. ISH for *neurog1* and *ascl1a* at the DC and SP stages in zebrafish showing that neural progenitors express the proneural *ascl1a* but not *neurog1*. Scale bar, 100µm **F**. Dorsal view (top panel) and cross section (bottom panel) of a 10dpf medaka pallium immunostained for GS (green) and Sox2 (blue), showing that the ventricular surface is paved with RG like in zebrafish. Scale bar, 50µm.

**Table S1:** List of primers used in this study to clone killifish genes

**Table S2:** List of secondary antibodies used in this study

