## supplemental data for "Mosaic heterochrony in neural progenitors sustains accelerated brain growth and neurogenesis in the juvenile killifish *N. furzeri*"

**Figure S4: Self-renewing and neurogenic AP maintain embryonic-like marker expression (related to figure 4)**

**A-A'.** Dorsal view of a killifish pallium at 13dph with an immunostaining for Sox2, GS and detection of EdU. A' is a higher magnification of the boxed area in A. Most EdU-labeled cells are AP. Scale bar, 50µm **B**. Proportion of EdU-labeled cells that are AP (blue, Sox2+GS-) or RG (green, Sox2+GS+) immediately following the EdU pulse (t0). Data are represented as mean ± SEM; n=2. **C**. Proportion of EdU-labeled AP cells that have incorporated BrdU immediately following the BrdU pulse (t1) or after 3 days of chase. Data are represented as mean ± SEM; n=2 (t1) or n=3(t2) for each treatment condition. Statistical analysis does not reveal any statistical difference (2-way ANOVA approach followed by pairwise comparisons, data were ranked transformed prior to analysis). **D**. *notch1* orthologs, *Dll1* orthologs and *neurog1* expression in the killifish pallium at 8dph. Whole-mount ISH of the pallium are shown as well as higher magnification in the top right corner. Some signal-positive cells are indicated by white arrows. A 50µm-vibratome section is also shown for *neurog1*, highlighting the salt-and-pepper expression in the ventricular layer. Scale bar, 100µm. **E**. ISH for *neurog1* and *ascl1a* at the DC and SP stages in zebrafish showing that neural progenitors express the proneural *ascl1a* but not *neurog1*. Scale bar, 100µm **F**. Dorsal view (top panel) and cross section (bottom panel) of a 10dpf medaka pallium immunostained for GS (green) and Sox2 (blue), showing that the ventricular surface is paved with RG like in zebrafish. Scale bar, 50µm.

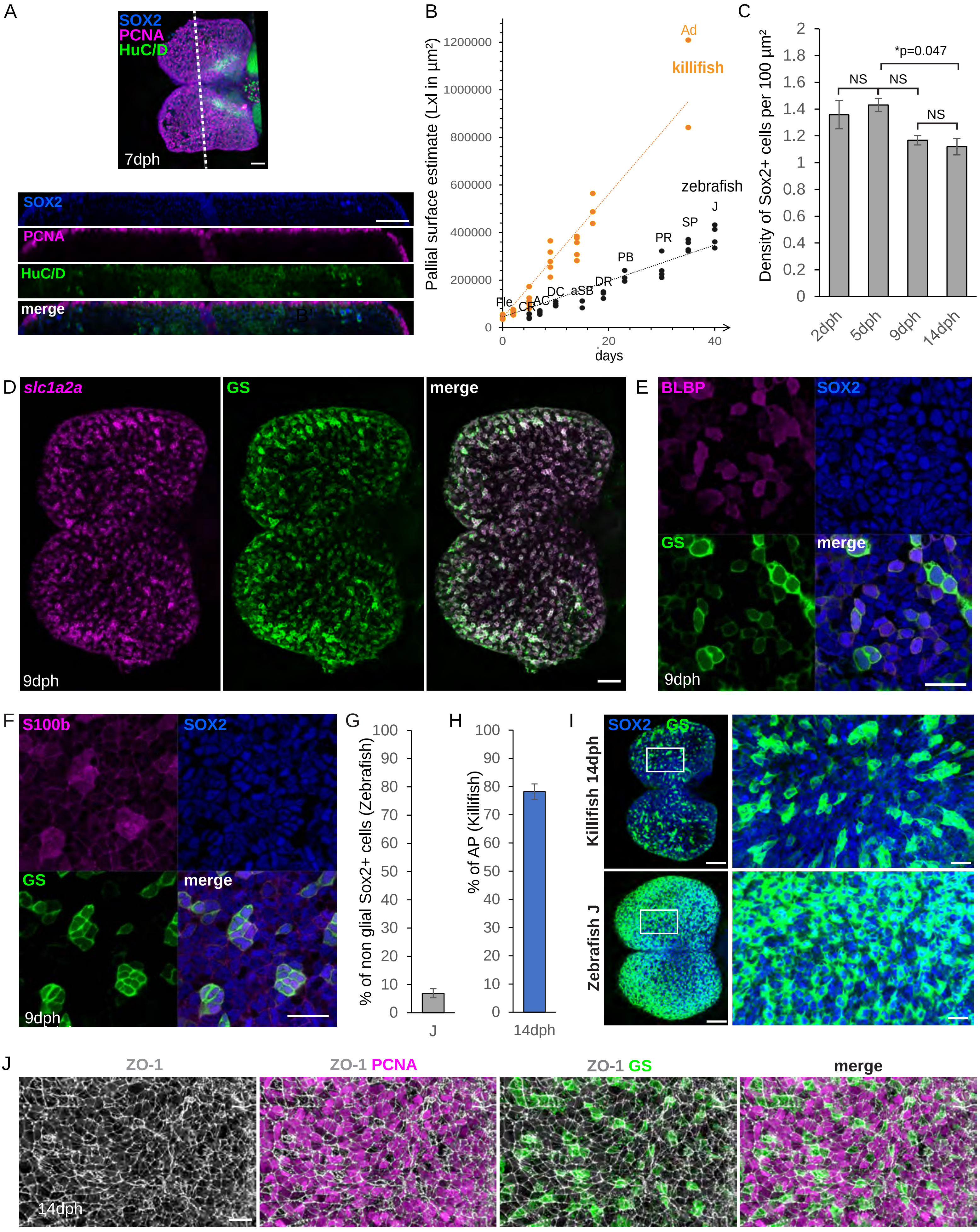
A
SOX2
PCNA
HuC/D
7dph
SOX2
PCNA
HuC/D
merge
B
Pallial surface estimate (LxI in µm²)
1200000
1000000
800000
600000
400000
200000
0
Ad
killifish
zebrafish
J
SP
PR
PB
DR
aSB
DC
AC
CR
Fle
0
20
40
days
C
Density of Sox2+ cells per 100 µm²
2
1.8
1.6
1.4
1.2
1
0.8
0.6
0.4
0.2
0
*p=0.047
NS
NS
NS
2dph
5dph
9dph
14dph
D
slc1a2a
GS
merge
9dph
E
BLBP
SOX2
GS
merge
9dph
F
S100b
SOX2
GS
merge
9dph
G
% of non glial Sox2+ cells (Zebrafish)
100
90
80
70
60
50
40
30
20
10
0
J
H
% of AP (Killifish)
100
90
80
70
60
50
40
30
20
10
0
14dph
I
SOX2 GS
Killifish 14dph
Zebrafish J
J
ZO-1
ZO-1 PCNA
ZO-1 GS
merge
14dph

Figure S1

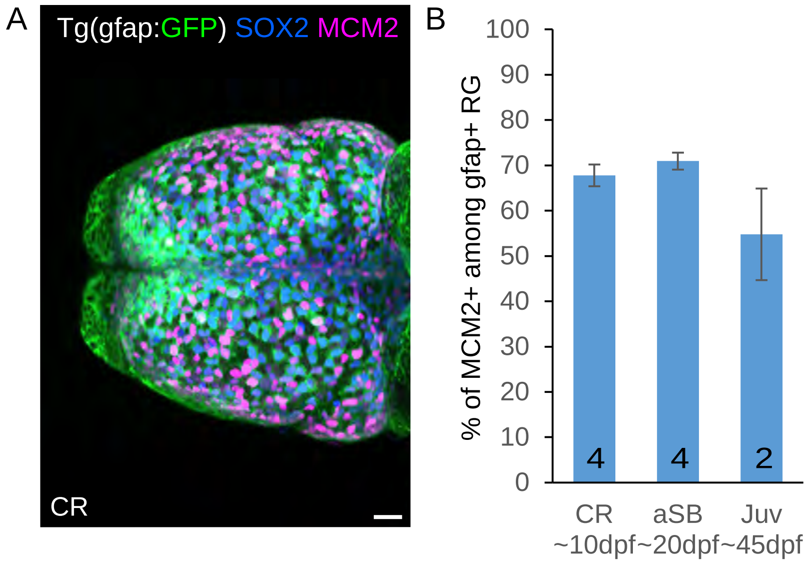
A
Tg(gfap:GFP) SOX2 MCM2
CR
B
% of MCM2+ among gfap+ RG
100
90
80
70
60
50
40
30
20
10
0
4
4
2
CR
~10dpf
aSB
~20dpf
Juv
~45dpf

Figure S2

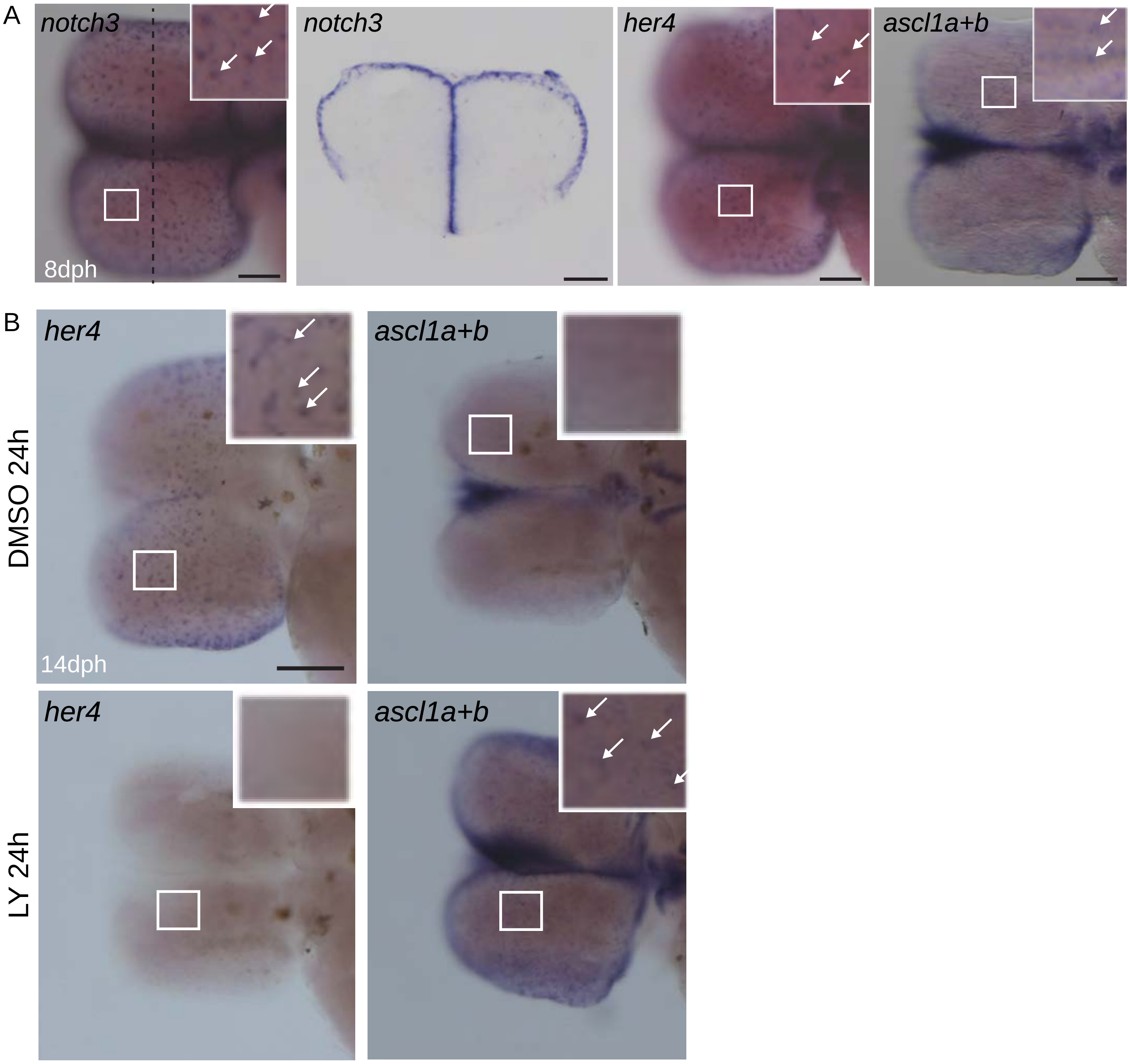
A
notch3
8dph
notch3
her4
ascl1a+b
B
her4
ascl1a+b
DMSO 24h
14dph
her4
ascl1a+b
LY 24h

Figure S3

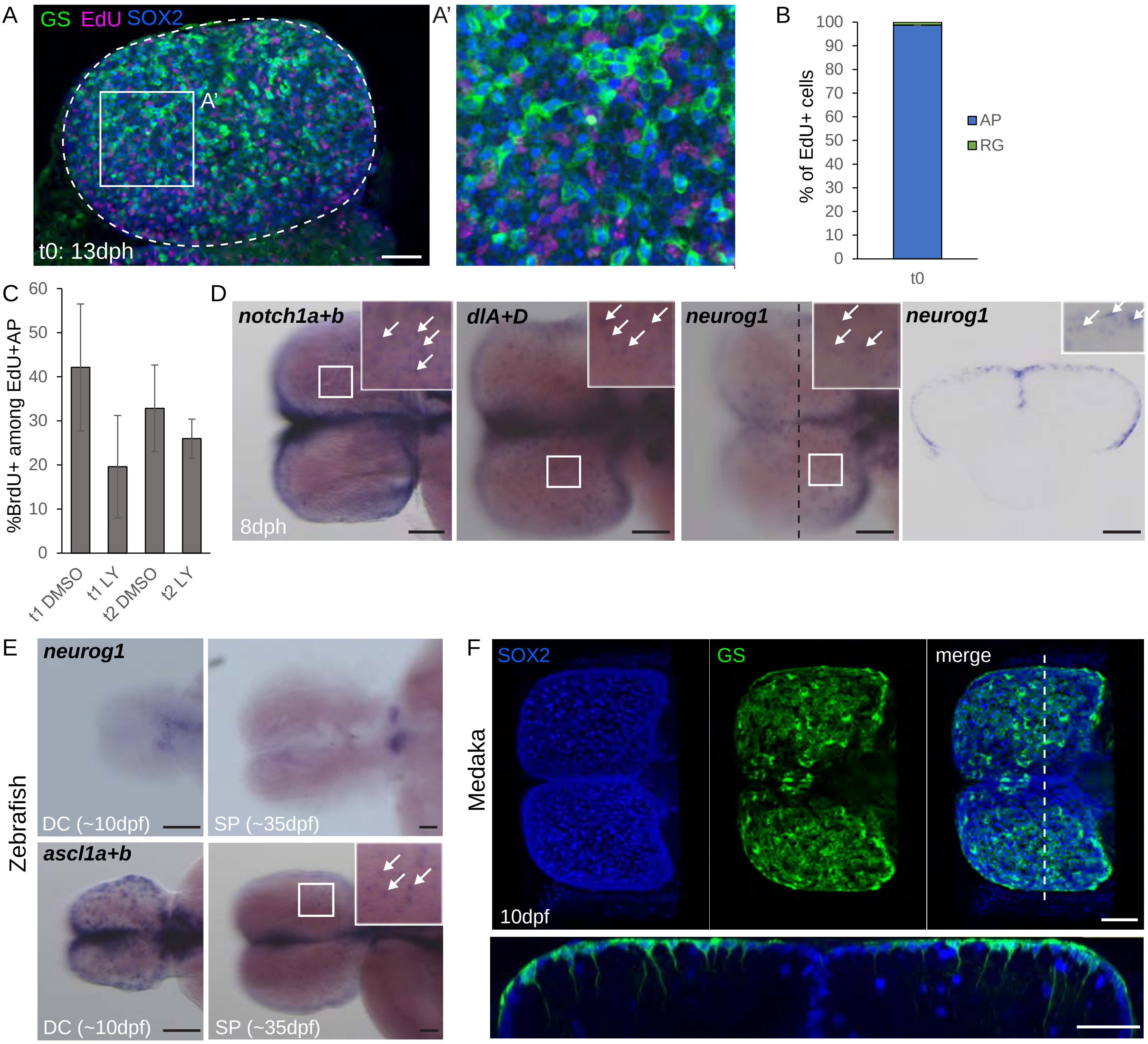
A
GS EdU SOX2
A'
t0: 13dph
A'
B
100
90
80
70
60
50
40
30
20
10
0
% of EdU+ cells
AP
RG
t0
C
%BrdU+ among EdU+AP
60
50
40
30
20
10
0
t1 DMSO
t1 LY
t2 DMSO
t2 LY
D
notch1a+b
dlA+D
neurog1
neurog1
8dph
E
Zebrafish
neurog1
DC (~10dpf)
SP (~35dpf)
ascl1a+b
DC (~10dpf)
SP (~35dpf)
F
Medaka
SOX2
GS
merge
10dpf

Figure S4

| Gene name | Genbank identifier | Zebrafish ortholog | Mouse ortholog | Primer FWD (5'-3') | Primer REV (5'-3') | insert length (bp) |
|---|---|---|---|---|---|---|
| ***Nf_ascl1a*** | GAIB01132435 | *ascl1a* | *Ascl1* | CCTGGACACTGGGTGACTTT | TCCGTGGTTCTGTGAGGA | 692 |
| ***Nf_ascl1b*** | GAIB01133487 | *ascl1b* | *Ascl1* | AAACTGCAACCATCACCACC | AGGGGCGGAAATGTACATCA | 995 |
| ***Nf_dlA*** | GAIB01004767 | *dlA* | *Dll1* | ATCTAATATGGGGCGCGTCA | CTCTCTCAGGGTTGTCAGCA | 1078 |
| ***Nf_dlD*** | GAIB01092115 | *dlD* | *Dll1* | GCATTCACAGTGACAGCCAA | CCATCCAAGTCATTCCTGCG | 1063 |
| ***Nf_her4.2*** | GAIB01137651 | *her4.2* | *Hes5* | ATGGCTCCTACAATCACTGC | CCGTAGATTCAGTCGTGCAA | 643 |
| ***Nf_neurog1*** | GAIB01038862 | *neurog1* | *Neurog1* | GAGCAAAGAAACGCATAGAGG | CCATCAGGAGGAGAAGTGGA | 975 |
| ***Nf_notch1a*** | GAIB01090429 | *notch1a* | *Notch1* | ACGGCTTGGCTTCCTTCA | GGGTTGCTCTCACATTCATT | 1114 |
| ***Nf_notch1b*** | GAIB01101509 | *notch1b* | *Notch1* | ACTGTGAGAGCCGTTACCTTCC | TACTTGTTTGGTCCGTCTGTG | 1008 |
| ***Nf_notch3*** | GAIB01090895 | *notch3* | *Notch3* | TTCCTAATCCCTGCCTAAATG | CCACCCACTCTATCCACACA | 1018 |
| ***Nf_slc1a2a*** | GAIB01110701 | *slc1a2a* | *slc1a* | CCAATCCATCCAGATATTGTCAT | CTTGCTACAACCTCCAAGTCCT | 707 |

**Table S1**: List of primers used in this study to clone killifish genes

| Secondary Antibody Name | Target species | Host species | Coupled to | Source | Identifier |
|---|---|---|---|---|---|
| DyLight 405-AffiniPure Goat Anti-Mouse IgG2a | Mouse IgG2a | Goat | DyLight405 | Jackson ImmunoResearch Labs | Cat#115-475-206, RRID:AB_2338800 |
| Goat anti-Mouse IgG2a-488 | Mouse IgG2a | Goat | Alexa Fluor 488 | Thermo Fisher Scientific | Cat#A-21131, RRID:AB_2535771 |
| Goat anti-Mouse IgG2a-546 | Mouse IgG2a | Goat | Alexa Fluor 546 | Thermo Fisher Scientific | Cat#A-21133, RRID:AB_2535772 |
| Goat anti-Mouse IgG2a-633 | Mouse IgG2a | Goat | Alexa Fluor 633 | Thermo Fisher Scientific | Cat#A-21136, RRID:AB_2535775 |
| Goat anti-Mouse IgG2b-633 | Mouse IgG2b | Goat | Alexa Fluor 633 | Thermo Fisher Scientific | Cat#A-21146, RRID:AB_2535782 |
| DyLight™ 405 Goat anti-Mouse IgG1 | Mouse IgG1 | Goat | DyLight405 | Biolegend | Cat#409109, RRID:AB_10642830 |
| Goat anti-Mouse IgG1-546 | Mouse IgG1 | Goat | Alexa Fluor 546 | Thermo Fisher Scientific | Cat#A-21123, RRID:AB_2535765 |
| Goat anti-Mouse IgG1-647 | Mouse IgG1 | Goat | Alexa Fluor 647 | Thermo Fisher Scientific | Cat#A-21240, RRID:AB_2535809 |
| Goat anti-Rabbit-546 | Rabbit IgG (H+L) | Goat | Alexa Fluor 546 | Thermo Fisher Scientific | Cat#A-11010, RRID:AB_2534077 |
| Goat anti-Rabbit-633 | Rabbit IgG (H+L) | Goat | Alexa Fluor 633 | Thermo Fisher Scientific | Cat#A-21071, RRID:AB_2535732 |
| Goat Anti-Chicken IgG-488 | Chicken IgY | Goat | Alexa Fluor 488 | Thermo Fisher Scientific | Cat#A-11039, RRID:AB_142924 |
| Goat anti-Rat IgG-488 | Rat IgG (H+L) | Goat | Alexa Fluor 488 | Thermo Fisher Scientific | Cat#A-11006, RRID:AB_2534074 |

**Table S2**: List of secondary antibodies used in this study
